# Molecular rationale for hantavirus neutralization by a reservoir host-derived monoclonal antibody

**DOI:** 10.1101/2020.04.17.029876

**Authors:** Ilona Rissanen, Robert Stass, Stefanie A. Krumm, Jeffrey Seow, Ruben J.G. Hulswit, Guido C. Paesen, Jussi Hepojoki, Olli Vapalahti, Åke Lundkvist, Olivier Reynard, Viktor Volchkov, Katie J. Doores, Juha T. Huiskonen, Thomas A. Bowden

**Author notes:** Equal contribution (Ilona Rissanen, Robert Stass).

## Abstract

The intricate lattice of Gn and Gc glycoprotein spike complexes at the surface of hantaviruses facilitates host-cell entry and is the primary target of the neutralizing antibody-mediated immune response. Here, through study of a neutralizing monoclonal antibody (mAb 4G2) generated in a bank vole reservoir host following infection with Puumala virus (PUUV), we provide molecular-level insights into how antibody-mediated targeting of the hantaviral glycoprotein lattice effectively neutralizes the virus. Crystallographic analysis reveals that mAb 4G2 binds to a multi-domain site on Gc in the pre-fusion state, and that Fab binding is incompatible with the conformational changes of the Gc that are required for host cell entry. Cryo-electron microscopy of PUUV-like particles treated with Fab 4G2 demonstrates that the antibody binds to monomeric Gc at breaks in the Gn-Gc lattice, highlighting the immunological accessibility of Gc monomers on the mature hantavirus surface and the plastic nature of the higher-order lattice assembly. This work provides a structure-based blueprint for rationalizing antibody-mediated targeting of hantaviruses.

## Introduction

Rodent-borne hantaviruses (genus *Orthohantavirus*, family *Hantaviridae*) are enveloped, negative-sense RNA viruses found worldwide in small mammals [1, 2]. Cross-species transmission into humans typically results from inhalation of aerosolized excreta from chronically infected rodents, and has two primary clinical outcomes. Hantaviral species native to Europe and Asia cause hemorrhagic fever with renal syndrome (HFRS), while those in the Americas cause hantavirus cardiopulmonary syndrome (HCPS) [1, 3, 4]. The case mortality rate may exceed 30 % for HCPS and ranges from <1% to 15% for HFRS [2, 5, 6]. Despite variable clinical characteristics, both syndromes arise from excessive proinflammatory and cellular immune responses to infection [7, 8]. Due to their potential to cause severe disease and to be transmitted by aerosols, hantaviruses have been identified as potential bioterrorism agents by the Centers for Disease Control and Prevention [9].

Entry of a hantavirus into a host-cell is negotiated by two membrane-anchored glycoproteins, Gn and Gc, which form a lattice decorating the lipid bilayer envelope of the mature virion [10–12]. Gn and Gc are translated as a single polypeptide, and cleaved at a ‘WAASA’ signal sequence during protein folding and assembly [13, 14]. Recent studies have reported crystal structures of both Gn [15, 16] and Gc [17, 18] ectodomains, and their higher-order, tetrameric organization has been postulated based upon biochemical characterization [10] and cryo-electron microscopy reconstructions of purified virions [11, 12, 16]. The Gn ectodomain exhibits a mixed α/β fold, which locates to the membrane-distal region of the tetrameric spike and is linked to the virion envelope by a C-terminal stalk region [11, 15, 16]. The Gc adopts a three-domain (domains I—III) class II fusion protein architecture, and is postulated to associate closely with the Gn, linking adjacent Gn tetramers [10, 16–18]. The Gn has been predicted to shield the fusion loop resident in domain II of the Gc, in a manner similar to that proposed in phleboviruses and alphaviruses [16, 17, 19–21]. Following host cell binding and endocytotic uptake of the virus, acidification in endosomal compartments results in at least partial dissolution of the Gn-Gc lattice [10, 15, 22]. Endosomal escape is mediated by the Gc in a process that involves engagement of the target membrane by the Gc-encoded fusion loop, followed by upward ‘zippering action’ conformational rearrangements of the transmembrane-anchored Gc domain III, which bring together host and viral membranes [17, 18].

Long co-evolution of hantaviruses with their reservoirs hosts has been proposed to have led to mechanisms by which the virus restrains host immune responses, including proinflammatory cytokine responses and CD8+ T-cell activity, thus allowing persistence of the virus in animal populations [23, 24]. This regulatory mechanism is likely defective in human infections, explaining the pathologies and efficient viral elimination observed in hantaviral disease [25]. A robust antibody-mediated immune response targeting Gn and Gc has been shown to be key in limiting disease progression upon zoonosis, and high titers of neutralizing antibodies (nAbs) in the acute phase correlate with a more benign course of disease [26–28]. A neutralizing humoral immune response also conveys long-lasting immunity against hantaviral infection [29–31]. Despite concerted efforts to understand the humoral immune response against pathogenic hantaviruses [32–35], little is known about the molecular determinants of antibody-mediated neutralization.

Here, we structurally characterize the epitope of mAb 4G2, a potently neutralizing antibody derived following experimental infection of bank voles with Puumala virus (PUUV) [32], a hantavirus responsible for one of the most frequently occurring hantaviral diseases, a mild form of HFRS known as nephropathia epidemica (NE) [6, 36]. Through a combined crystallographic and electron cryo-tomography (cryo-ET) analysis, we report the structure of mAb 4G2 in complex with PUUV Gc and define the 4G2 epitope on the Gc subcomponent of the mature Gn-Gc spike. This work provides an initial molecular-level basis for understanding immune-mediated targeting of the antigenic hantaviral surface.

## Results

### Recombinantly-derived and Gc-specific bank vole mAb 4G2 potently neutralizes PUUV

The hybridoma cell line producing the Gc-specific neutralizing mAb 4G2 was generated in 1992 following experimental infection of the natural reservoir species, bank voles (*Myodes glareolus*) with PUUV [32]. While mAb 4G2 has been shown to be potently neutralizing against PUUV, with a mAb concentration of ~4 nM (0.6 ug/ml) resulting in 80% inhibition of PUUV infection in focus reduction neutralization tests [32, 37, 38], there has been a paucity of information with regards to the structural basis for neutralization by this or any other Gc-specific antibody. We rescued the sequence of the 4G2 variable heavy (V_H_) and kappa (V_K_) chain regions (see *Supporting information,* **Fig S1**) from hybridoma cells by PCR with gene specific primers originally designed for mouse V_H_ and V_K_ chains [39]. The resulting recombinantly-produced mAb 4G2 comprised bank vole variable and mouse constant regions and, consistent with previously reported *in vitro* studies of hybridoma-derived mAb 4G2 [10, 32], potently neutralizes PUUV in a hantavirus-pseudotyped VSV-ΔG RFP neutralization assay (**Fig 1**).

**Figure 1.**
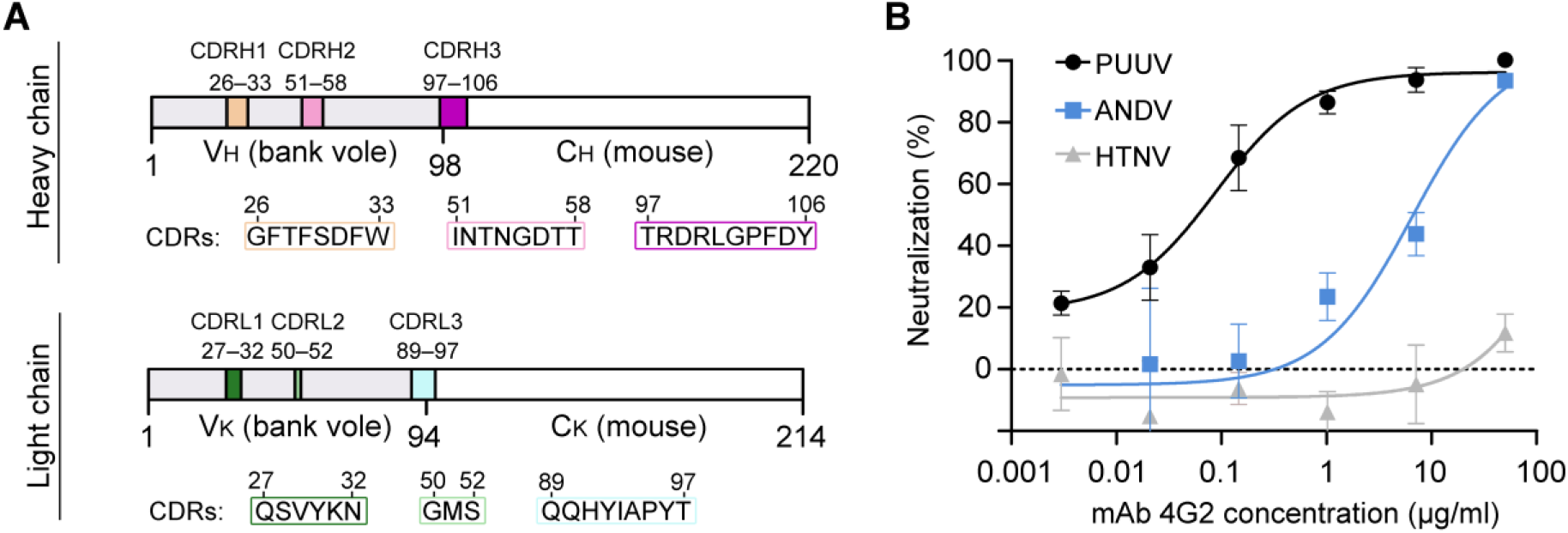
Composition and neutralization potency of rescued bank vole mAb 4G2. (**A**) Composition of the complementarity determining regions (CDRs) of the mAb 4G2 antigen binding fragment (Fab) heavy (VH) and kappa (VK) chains. For the full sequence of Fab 4G2 variable regions, please see *Supporting information*, **Fig S1**. (**B**) A hantavirus-pseudotyped VSV-ΔG RFP neutralization assay shows that recombinantly-produced mAb 4G2 neutralizes PUUV- and ANDV-pseudotyped VSV (black and blue traces, respectively), but not HTNV-pseudotyped VSV (grey trace). Each neutralization assay was carried out three times in duplicate. A representative experiment is shown. Error bars represent the range of the value for the experiment performed in duplicate.

### Crystal structure of the Fab 4G2 in complex with PUUV Gc reveals the 4G2 epitope on pre-fusion Gc

To ascertain the molecular basis for 4G2-mediated recognition of PUUV, we determined the crystal structure of the PUUV Gc ectodomain in complex with the antigen-binding fragment (Fab) of 4G2 to 3.50-Å resolution (**Fig 2**, and *Supporting information* **Table S1** and **Fig S2**). Consistent with previous studies of hantaviral Gc glycoproteins [17, 18], PUUV Gc assumes a class II fusion glycoprotein architecture composed of three domains (domains I—III), with domain I and domain II forming an elongated structure and domain III forming inter-domain contacts at the side of domain I. Fab 4G2 recognizes a protein-specific epitope at the junction of domain I and domain II of PUUV Gc, distal from the hydrophobic fusion loop at the tip of domain II (**Fig 2**). The epitope comprises ~830 Å^2^ of buried surface area and all complementarity-determining region (CDR) loops participate in the interaction (**Fig 2B**). Arg100 (CDRH3) is the most centrally located paratope residue, and forms a salt bridge with Gc Glu725 and multiple hydrogen bonds across the antibody-antigen interface. Indeed, site-directed mutagenesis of Arg100 to Ala considerably reduced the potency of 4G2 to neutralize PUUV (*Supporting information*, **Fig S3**). One Fab 4G2-PUUV Gc complex was observed in the crystallographic asymmetric unit, and there was no evidence for the formation of higher-order Gc dimers or trimers, such as those observed in pre- and post-fusion class II fusion glycoprotein structures [40].

**Figure 2.**
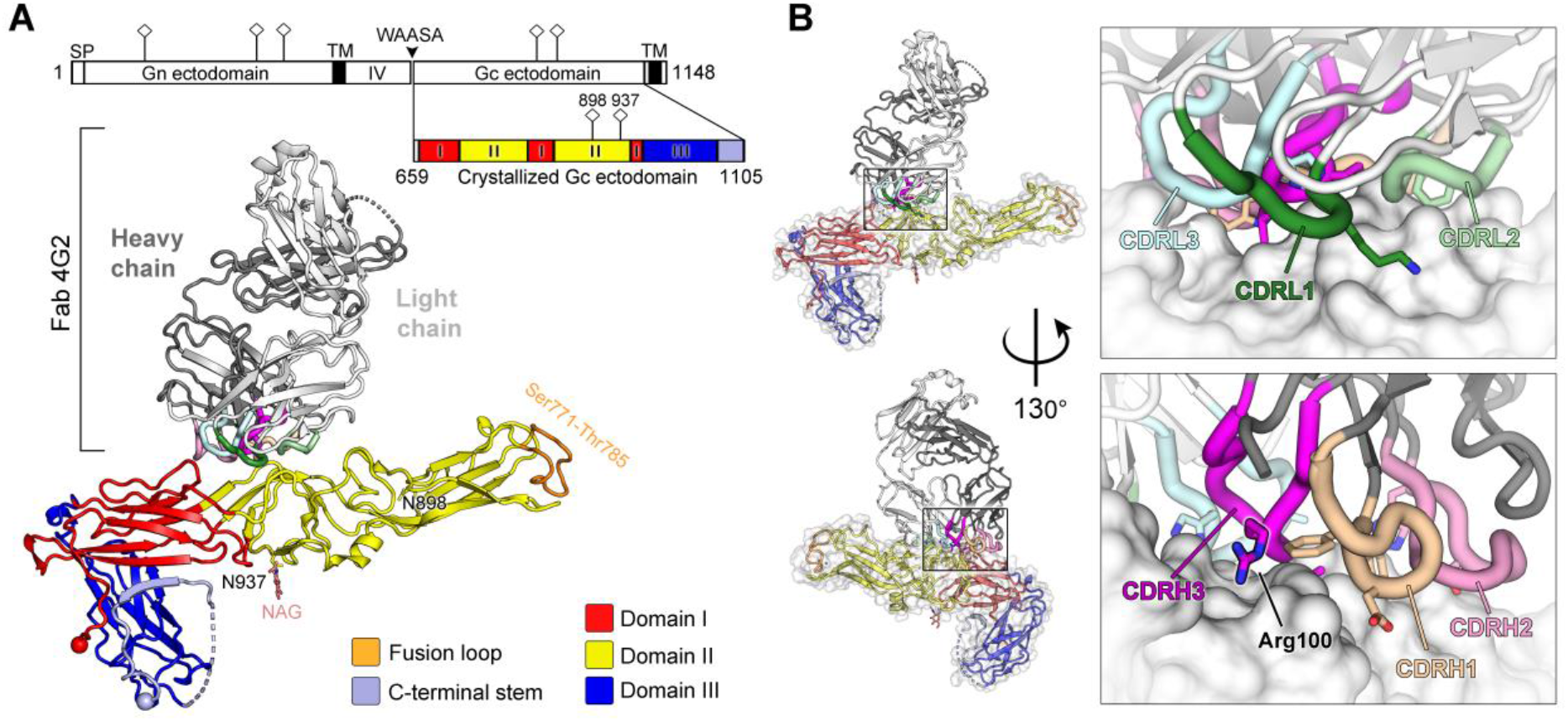
Crystal structure of neutralizing antibody 4G2 in complex with PUUV Gc. (**A**) Crystal structure of Fab 4G2-PUUV Gc complex at 3.5 Å. PUUV Gc, a class II fusion protein, comprises domains I-III (colored red, yellow, and blue, respectively), a Gc C-terminal tail (light blue) and the viral fusion loop at Ser771-Thr785 (orange). Fab 4G2, comprised of a heavy chain (dark grey) and a light chain (white), is observed bound at the junction of domains I and II on PUUV Gc. Domain schematic of the PUUV glycoprotein precursor with the signal peptide (SP), transmembrane domains (TM), intra-virion domain (IV), and WAASA signal peptidase cleavage site is shown alongside the construct used in crystallization (schematic was produced using the DOG software [76]). N-linked glycosylation sequons are shown in the domain schematic as pins, and sites at Asn898 and Asn937 are highlighted in the structure representation, where the *N*-linked glycan observed at Asn937 is shown as sticks. There was no evidence of glycosylation at Asn898. (**B**) The interface of Fab 4G2 with PUUV Gc. CDRs contributing to PUUV recognition are highlighted and key buried residues, identified by the PDBePISA server [77], are rendered at sticks, including the centrally located CRDH3 Arg100.

Consistent with the hypothesis that mAb 4G2 targets a pre-fusion conformation of PUUV Gc representative of that displayed on the native virion surface, structure overlay analysis reveals that PUUV Gc from the Gc-4G2 complex (PUUV Gc_Gc-4G2_) adopts a conformation distinct from the known post-fusion structure [18]. Furthermore, the conformation of PUUV Gc_Gc-4G2_ closely resembles class II fusion protein pre-fusion states, such as those observed for RVFV Gc (PDB 4HJ1) and ZIKV E (PDB 5IRE) (*Supporting information*, **Fig S4**). Indeed, an investigation into the residues that comprise the 4G2 epitope revealed that these residues are buried in the post-fusion trimeric Gc (*Supporting information*, **Fig S5**), indicating that mAb 4G2 is specific to the pre-fusion state of PUUV Gc. This observation is consistent with previous experimental findings, which demonstrated that mAb 4G2 did not recognize PUUV under low pH conditions but remained bound to the virus and protected the epitope if introduced prior to low-pH exposure [10]. Taken together, these results support a model whereby 4G2 specifically targets pre-fusion Gc and may sterically preclude the formation of a fusogenic configuration of the Gc.

### The epitope of 4G2 is targeted for neutralization across hantaviral species

Hantaviruses are often divided into two groups, termed Old World and New World hantaviruses [2], reflecting their distribution and pathobiological features. Interestingly, while PUUV is an Old World hantavirus prevalent in Northeastern Europe, the residues comprising the 4G2 epitope on PUUV Gc exhibit a relatively high level of sequence conservation with Gc proteins from New World hantaviruses, such as ANDV (72% sequence identity, with only nine epitope residues differing between the two proteins) (*Supporting information*, **Fig S5**). The observed similarity at the epitope between PUUV and New World hantaviruses is in line with the evolutionary history of these viruses and their hosts [41], and provides a structural basis for the ability of 4G2 to weakly neutralize ANDV (**Fig 1B**). Furthermore, despite the sequence variation at the 4G2 epitope between the two Old World hantaviruses PUUV and HTNV (**Fig 3**), and the lack of HTNV cross-neutralization (**Fig 1B**), we note that the observed 4G2 binding site overlaps with the putatively assigned binding sites of HTNV-neutralizing mAbs HCO2 and 16E6 [33, 42] (**Fig 3**), indicating that this region of the Gc is likely to be immunologically accessible across hantaviral species.

**Figure 3.**
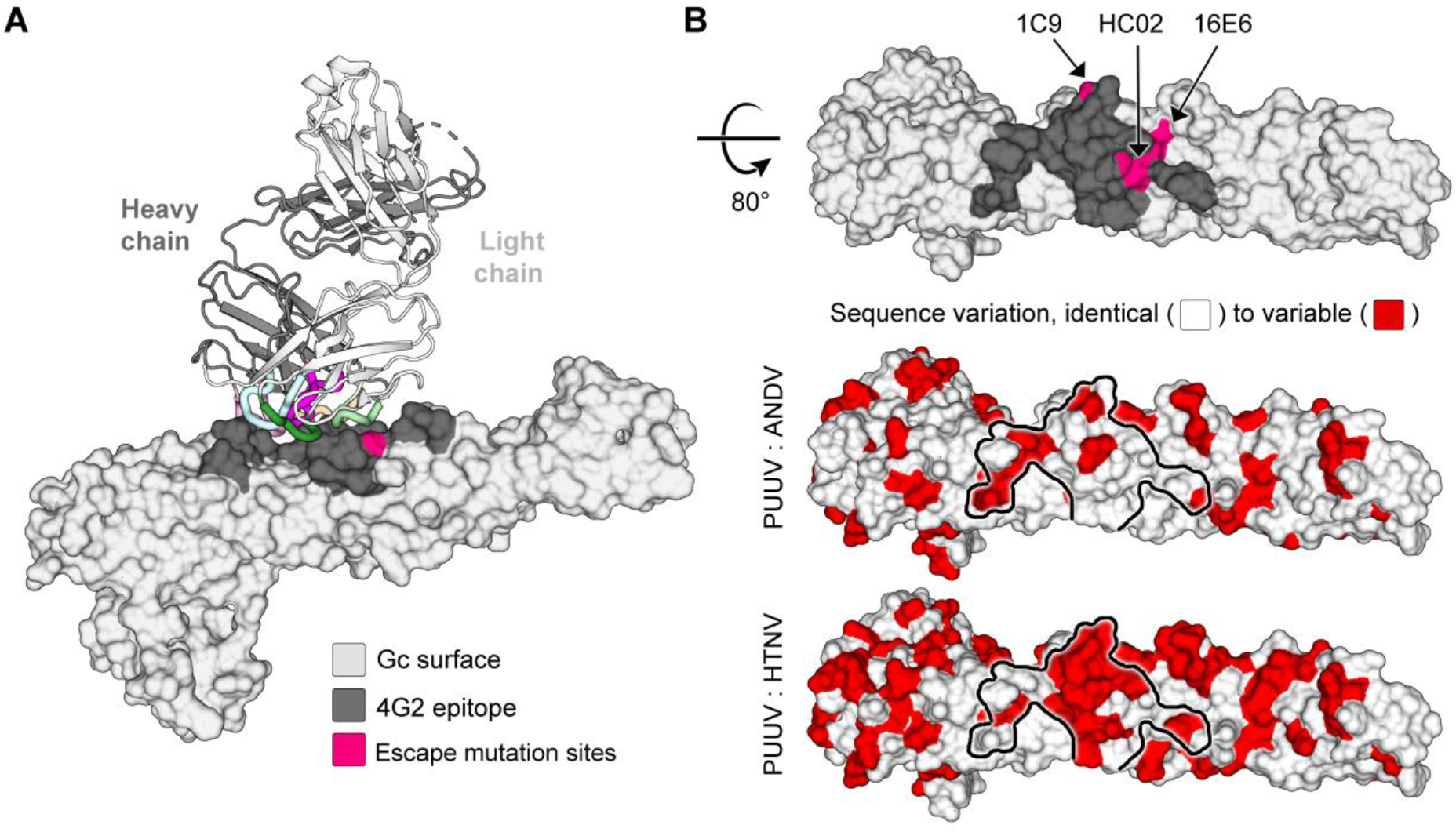
The epitope of antibody 4G2 denotes a key antigenic site at the hantaviral surface. (**A**) The epitope of 4G2 (dark grey, interfacing residues identified by the PDBePISA server [77]) is shown on the surface of PUUV Gc (light grey) and overlaps with the neutralization evasion mutation sites reported for PUUV-neutralizing human antibody 1C9 (S944F) [48, 49] and HTNV-neutralizing mouse antibodies HC02 and 16E6 (K795Q and L719P) [33, 42]. (**B**) Top view of the epitope, along with pairwise comparisons of PUUV Gc sequence to the sequences of Gc from ANDV and HTNV. For each pairwise comparison, the surface of PUUV Gc is shown colored white for identical residues and red for variable residues. The relatively high level of similarity between PUUV and ANDV is in line with the observed, albeit weak, neutralization of mAb 4G2 against ANDV (**Fig 1**).

### Mab 4G2 is specific to a monomeric state of virion-displayed PUUV Gc

Previous investigations of the non-pathogenic viral orthologue, Tula virus (TULV), have revealed the ultrastructure of Gn-Gc assemblies, as displayed on the hantaviral envelope [11, 16]. These studies indicate that the Gc forms elongated structures that extend from the virion membrane and are shielded by the cognate Gn [16].

To ascertain the position of the 4G2 epitope in the context of the hantaviral envelope, we generated PUUV virus-like particles (VLPs) by transient expression of the PUUV genomic M-segment in mammalian cells [43]. Cryo-electron tomography (cryo-ET) of purified PUUV VLPs, in tandem with sub-tomogram averaging, revealed that the VLP surface displays the expected Gn-Gc spike architecture that can form locally ordered lattices, as observed in native hantavirions (*Supporting information*, **Fig S6**). Consistent with previous studies of hantavirus ultrastructure and Gc oligomerization [16, 44], breakpoints presenting monomeric Gc exist in the (Gn-Gc)_4_ lattices [11, 16].

Following the validation that PUUV VLP accurately resembles the native virion, we applied an identical cryo-ET approach to VLPs treated with Fab 4G2. The data were split in two sets: (i) spikes that were part of a lattice, and (ii) spikes in regions of incomplete lattice. The former yielded a reconstruction similar to the VLP sample prepared without the Fab (**Fig 4A** and *Supporting information*, **Fig S6**), while the reconstruction focused on lattice-free spikes yielded a discrete (Gn—Gc)_4_ spike with additional density (**Fig 4A**). These results suggest that mAb 4G2 binding is incompatible with the metastable Gc homodimerization interface, and further, that Fab 4G2 binding may affect the higher order lattice assembly. To test this hypothesis, we quantified the number of neighbors for each spike on VLPs both in the presence and absence of Fab 4G2. The frequency of lattice-free spikes (with zero neighbors) was higher in the presence of Fab 4G2, while the frequency of lattice-bound spikes (with four or more neighbors) was higher in the particles not treated with Fab 4G2 (**Fig 4B** and *Supporting information*, **Fig S7**). These results suggest that Fab 4G2 influences the presentation of the Gn—Gc lattice by dislodging the (Gn—Gc)_4_ spikes from each other.

**Figure 4.**
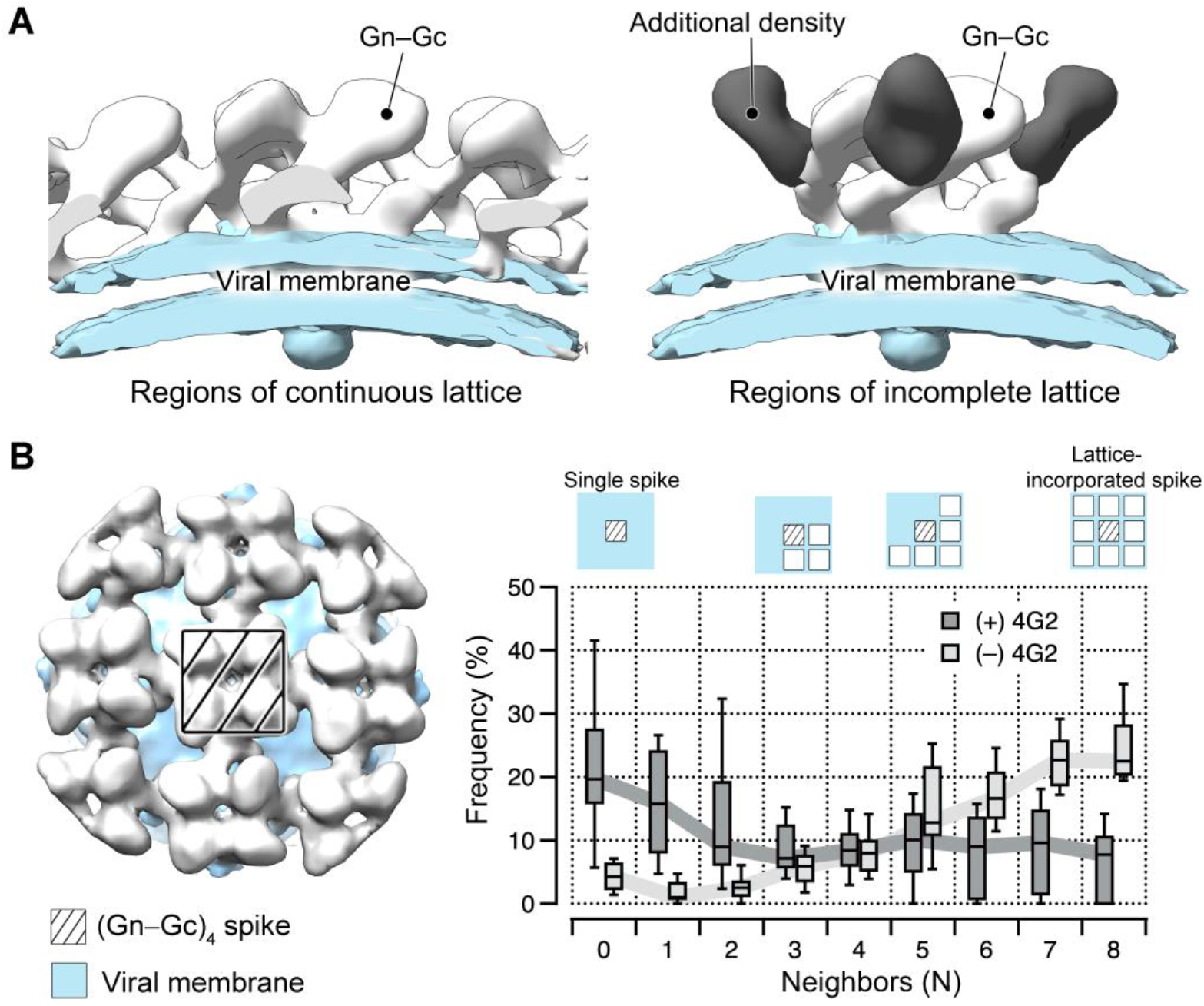
Treatment of PUUV VLPs with Fab 4G2 results in additional density and is associated with loss of continuous lattice at the VLP surface. (**A**) Two cryo-ET reconstructions of Fab 4G2-treated PUUV VLP surface, derived from (i) regions of continuous lattice (16.8 Å) and (ii) regions of incomplete lattice (15.0 Å). While both reconstructions show canonical Gn-Gc architecture (density colored white) and the viral lipid bilayer (light blue), additional density is observed in the latter reconstruction (dark grey). (**B**) The hantaviral surface carries tetragonal (Gn-Gc)_4_ spikes that can organize in patches of ordered lattice. A box plot describing the frequency of (Gn-Gc)_4_ spikes that have a given number of lattice compatible neighbors, from zero to a maximum of eight, shows that treatment with Fab 4G2 alters the presentation of the (Gn-Gc)_4_ spike assemblies at the VLP surface.

Fitting of the crystal structure of Fab 4G2—PUUV Gc complex into the reconstruction of the (Gn—Gc)_4_ spike confirms that the additional density is comprised of Fab 4G2 (**Fig 5**). We also note that PUUV Gc_Gc—4G2_ fits well within the cryo-ET reconstruction of the spike (**Fig 5**), further supporting that PUUV Gc has crystallized in the pre-fusion conformation presented on the mature virion. Additionally, this fitting confirms the location of Gc in the density spanning from the membrane-distal globular lobes to the viral membrane, as we have proposed previously [16]. The fitting locates domain III of the Gc adjacent to the membrane, allowing the transmembrane region of the C-terminus to be linked to the virion envelope (**Fig 5B, Fig 2A**), and places the fusion loops from Gc domain II into membrane-distal lobe density in close contact with Gn (**Fig 5B**), in a manner similar to that observed for other class II fusion proteins and their cognate accessory proteins [20, 21]. Altogether, our combined crystallographic and cryo-EM investigation reveals, in molecular detail, that 4G2 is specific to the monomeric, pre-fusion state of PUUV Gc presented on the mature viral envelope.

**Figure 5.**
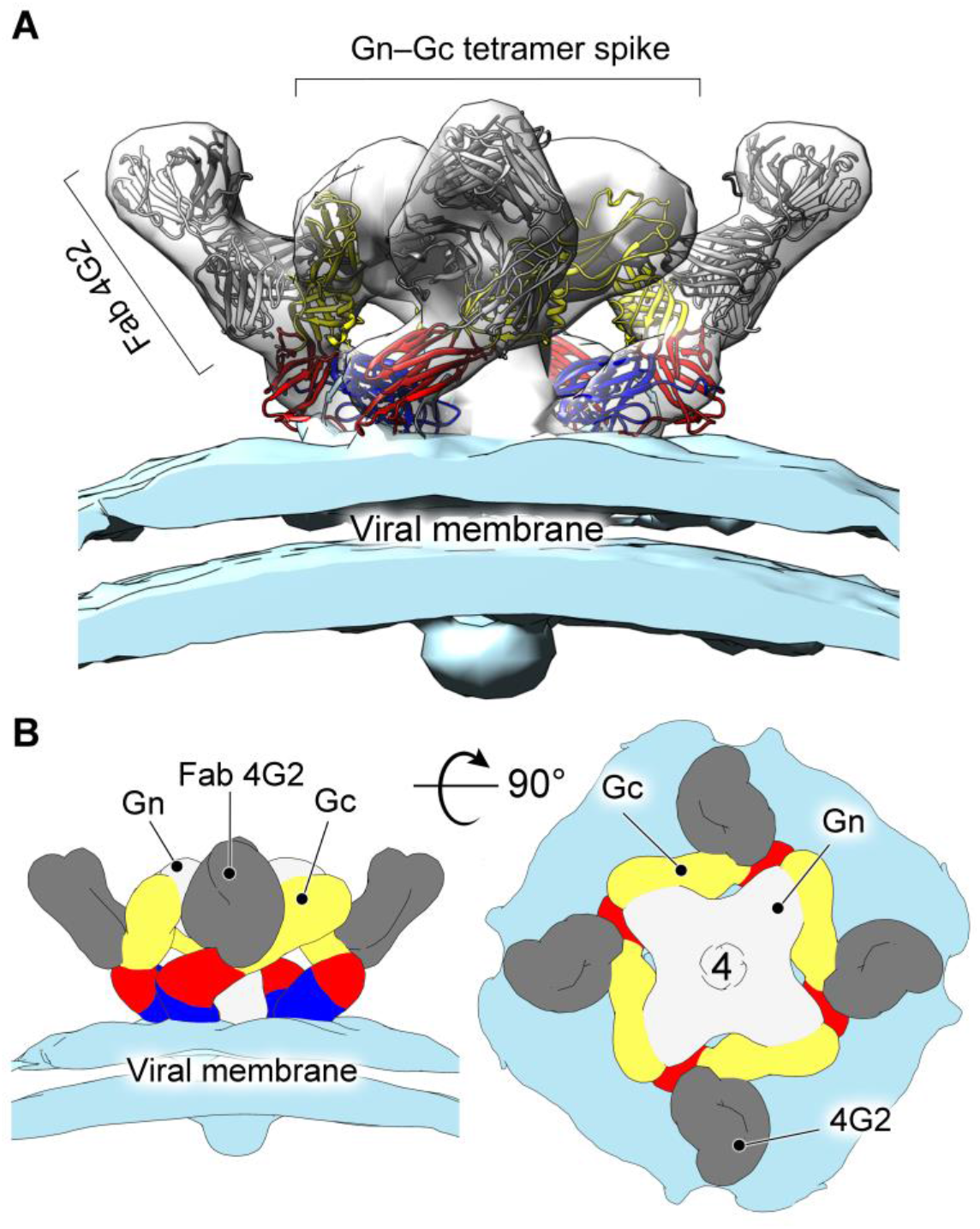
Fitting of the of Fab 4G2-PUUV Gc crystal structure into the cryo-ET reconstruction confirms the 4G2 epitope in the context of the viral surface and supports that the observed Gc conformation constitutes a pre-fusion state (**A**) Side view of Fab 4G2-treated PUUV VLP spike at 15-Å resolution. The crystal structure of Fab 4G2-PUUV was fitted in the cryo-ET reconstruction as a rigid body and displays excellent conformity with the cryo-ET derived envelope. PUUV Gc and Fab 4G2 heavy and light chains are colored as in **Fig 2**, and the lipid bilayer that constitutes the viral membrane is colored light blue. (**B**) Schematic representation of the organization of Gn_4_-Gc_4_ glycoprotein spike with Fab 4G2 bound.

## Discussion

Despite the importance of neutralizing antibodies in mitigating the progression of hantaviral disease [26–28] and conveying long-lasting protection against infection [31, 45, 46], little is known about how they target virion-displayed glycoproteins, Gn and Gc. Here, we provide a molecular-level description of the epitope targeted by mAb 4G2, a host-derived antibody that potently neutralizes PUUV, a zoonotic hantavirus prevalent in Northern Europe and Russia. We find that mAb 4G2 targets an epitope at the junction of domains I and II of PUUV Gc (**Fig 2**). When combined with a cryo-ET reconstruction of Fab 4G2 complexed with PUUV (Gn—Gc)_4_ spike, our analysis confirms previous postulations about the organization of the spikes on the hantavirus envelope [10, 16], and indicates that mAb 4G2 targets a pre-fusion, monomeric conformation of PUUV Gc.

Examination of the antibody—antigen interface provides a structure-based model for understanding the mechanism of 4G2-mediated neutralization, where overlay analysis reveals that Fab 4G2 binding is incompatible with the pH-dependent conformational transitions of PUUV Gc required to negotiate fusion of the viral and cellular membranes. While it is also possible that mAb 4G2 interferes with other steps of the host-cell entry pathway (e.g. receptor recognition), our modeling indicates that the antibody targets residues crucial for the formation of a post-fusion Gc configuration (*Supporting information*, **Fig S5**). Indeed, binding to this site likely impedes the upwards-occurring ‘zippering action’ of domain III during Gc-mediated viral fusion [17, 18]. Consistent with this proposed structure-based mechanism of neutralization, previous reports have shown that hindering such movements of domain III, either via mutagenesis [18] or the introduction of competing peptides [47], drastically reduces viral infectivity. Further supporting this model, mAb 4G2 binding has been reported to protect the epitope upon low-pH exposure [10].

In line with the hypothesis that the identified epitope on Gc domains I and II may constitute a site of vulnerability across hantaviruses, more generally, structure-based mapping revealed that the 4G2 binding site overlaps with epitopes putatively targeted by neutralizing antibodies specific to HTNV and PUUV. These antibodies, derived following hantaviral infection, include mouse mAbs HC02 and 16E6 [33, 42] that neutralize HTNV, and human mAb 1C9 [48, 49], which neutralizes PUUV (**Fig 3**). Thus, while our identified epitope appears to be a common target of the neutralizing antibody response, future investigations into the molecular basis of immune recognition will undoubtedly be required to fully define the antigenic topography of Gn and Gc. Such investigations will also likely clarify whether the neutralizing epitope defined here is similarly targeted upon immunization, and whether the Ab-immune response is differentially focused against Gn and Gc between reservoir species (e.g. bank vole) and accidental (i.e. humans) hosts.

Both dimers and monomers of Gc are expected to transiently exist on the hantavirus envelope: contacts between (Gn-Gc)_4_ spikes are mediated by Gc dimers, while lattice breaks display Gc monomers [11, 16, 44]. In our cryo-ET reconstruction, Fab 4G2 was found to bind PUUV Gc only in the monomeric state (*i.e*. Gc molecules that are not integrated within the lattice assembly) and the frequency of lattice-free spikes was higher in the presence of Fab 4G2. This observation demonstrates that 4G2 reduces the frequency of Gc-mediated lattice formation. While it is likely that the bivalent binding mode of full-length mAb 4G2, with respect to the structurally characterized Fab 4G2, may result in a different extent of lattice deformation, it is likely that both the mAb and Fab function by either active destabilization of Gc-Gc interfaces, or by sequestration of naturally occurring free spikes in a process that drives the equilibrium between dimeric and monomeric Gc towards the monomeric state.

The identification of the Gc oligomerization-specific epitope has important implications for rational therapeutic design efforts and indicates that Gc monomers can be specifically targeted by the antibody-mediated immune response that arises during infection. Indeed, although the development of recombinant immunogens that recreate the higher-order hantaviral (Gn-Gc)_4_ lattice is desirable and would likely elicit a robust immune response, our data also indicate that a reverse vaccinology approach [50] focusing on recombinant and more minimal (monomeric) hantaviral Gn or Gc may also elicit potent anti-hantaviral nAbs.

At present, treatment and prevention options for hantaviral disease remain extremely limited due to the absence of approved therapeutics. However, strides are being made towards the development of efficacious treatments, as shown by the recent development of novel mAbs against ANDV [51], and a polyclonal antibody treatment against HTNV [52], both of which demonstrate efficacy in animal models. *In toto*, our integrated structural and functional analysis provides a structure-based blueprint for understanding antibody-mediated neutralization of this group of emerging pathogens, and a template for the rational development of much needed biologics against this group of medically important pathogens.

## Materials and Methods

### Sequencing mAb 4G2 variable regions from hybridoma cell line

RNA from the 4G2 hybridoma cell line was converted into cDNA (SuperScript III reverse transcriptase, Life Technologies) using random hexamers following the manufacturer’s protocol. The bank vole antibody variable regions of heavy and kappa chains were PCR amplified using previously described mouse primers and PCR conditions [39]. PCR products were purified and cloned into mouse IgG expression plasmids [39] using sequence and ligation independent cloning (SLIC) under ampicillin selection. Antibody variable regions were sequenced by Sanger sequencing, and CDRs were determined from the sequences with IMGT/V-QUEST [53] server using *Mus musculus* as comparison group.

Antibody heavy and light plasmids were co-transfected at a 1:1 ratio into HEK293F cells (Thermo Fisher Scientific) using PEI Max 40K (linear polyethylenimine hydrochloride, Polysciences, Inc.), as previously described [54]. Antibody supernatants were harvested seven days following transfection and purified using protein G affinity chromatography following the manufacturer’s protocol (GE healthcare). In order to validate the functional relevance of Arg100, the R100A mutation was introduced into the 4G2 heavy chain plasmid using site directed mutagenesis (Forward primer: GTATTACTGTACAAGAGATGcATTAGGcCCTTTTGA, Reverse primer: TCAAAAGG GcCTAATGcATCTCTTGTACAGTAATAC) and the mutation was verified using Sanger sequencing. Mab 4G2 R100A mutant was expressed and purified as described above for WT mAb 4G2.

### Generation of hantavirus-pseudotyped VSV-ΔG RFP

8 × 10^6^ HEK 293T cells were transfected with a plasmid encoding PUUV GPC (35 μg in pCAGGS vector) and Tim-1 (3 μg) using PEI (3:1 PEI:DNA, Polysciences, Inc.). After 24 h, cells were infected with a VSV-G pseudotyped VSV-ΔG-RFP stock (40 uL). Infection was monitored using RFP expression and virus (1^st^ stock) was harvested when cells showed rounding and began to detach (typically 24-48 h).

Two additional rounds of transfection and virus production were carried out. 15 × 10^6^ HEK 293T cells were transfected with a plasmid encoding PUUV GPC, ANDV GPC or HTNV GPC (100 μg, in pCAGGS vector) and Tim-1 (6 μg) using PEI (3:1 PEI: DNA, Polysciences, Inc.). After 24 h, cells were infected with 400 μL of ’1^st^ stock’ virus. Infection was monitored using RFP expression and virus was harvested (2^nd^ stock) when cells showed rounding and began to detach (48-72 h). This process was repeated to generate ’3^rd^ stock’ virus which was used in subsequent neutralization assays.

### Hantavirus-pseudotyped VSV neutralization assay

Neutralizing activity was assessed using a single-round replication pseudovirus assay with HEK 293T target cells. Briefly, the antibody was serially diluted in a 96-well black, flat-bottom plate and preincubated with virus for 1 h at 37 °C. Cells at a concentration of 30,000/well were added to the virus-antibody mixture, and RFP quantified 72 h following infection (Envision plate reader, Perkin Elmer). Dose-response curves were fitted using nonlinear regression (GraphPad Prism) to determine 50% inhibitory concentration (IC_50_).

### Recombinant protein expression and purification

Codon-optimized synthetic cDNA (GeneArt, Life Technologies) coding for Fab 4G2 light chain and Fab 4G2 Heavy chain were individually cloned into the pHLsec [55] mammalian expression vector and co-transfected into human embryonic kidney (HEK) 293T cells (ATCC CRL-1573) for transient expression, as previously described [56]. Fab 4G2-containing supernatant was collected, clarified by centrifugation, and diafiltrated using the ÄKTA Flux tangential flow filtration system. Diafiltrated cell supernatant was purified by immobilized nickel affinity chromatography (5-ml fast flow crude column and ÄKTA fast protein liquid chromatography [FPLC] system; GE Healthcare) at room temperature, using 250mM imidazole for elution. Finally, the sample was purified by size exclusion chromatography (SEC) using a Superdex 200 10/300 Increase column (GE Healthcare), in 10 mM Tris (pH 8.0)-150 mM NaCl buffer.

In order to produce PUUV Gc, a HEK293T cell line stably expressing the protein was generated. PUUV Gc residues 659-1105 (GenBank accession no. CAB43026.1), modified to contain a 3C protease cleavable SUMO tag and hexahistidine tag at the N-terminus and a 1D4 tag in the C-terminus [57], were cloned into the pURD vector [58]. The stable cell line was generated following the method published by Seiradake et al [59]; in short, HEK293T cells were transfected with the pURD vector containing the Gc insert, along with an integrase expression vector (pgk-φC31/pCB92) [60], followed by consecutive rounds of cultivation under puromycin selection. For crystallization, PUUV Gc-producing cells were cultivated in the presence of kifunensine [61], and purification was performed via successive diafiltration, nickel affinity and size exclusion chromatography, as described above for Fab 4G2. Oligomannose glycans derived from expression in the presence of kifunensine were retained in the purified Gc sample.

Prior to crystallization, the N-terminal SUMO and hexahistidine tags were cleaved from the PUUV Gc (1:10 molar ratio of protein to 3C protease added to purified sample, followed by incubation at 21 °C for 12 hours). PUUV Gc sample was then mixed with pure Fab 4G2 in 1:1.2 molar ratio, and the complex was purified by SEC using a Superdex 200 10/300 Increase column (GE Healthcare), in 10 mM Tris (pH 8.0)-150 mM NaCl buffer.

### Crystallization and structure determination

A solution containing a complex of PUUV Gc ectodomain and Fab 4G2 was crystallized by sitting drop vapor diffusion using 100 nl protein (1.3 mg/mL) with 100 nl precipitant (20% vol/vol PEG 6000, 0.1 M MES pH 6 and 1 M lithium chloride) and 100 nl additive (6% 2-methyl-2,4-pentanediol) at room temperature. Crystals formed over two weeks and were flash frozen by immersion into a cryo-protectant containing 25% (vol/vol) glycerol followed by rapid transfer into liquid nitrogen. X-ray diffraction data were recorded at beamline I24 (λ = 0.9686 Å), Diamond Light Source, United Kingdom.

Data were indexed and integrated with XIA2/XDS [62] and scaled using CCP4/SCALEPACK2MTZ [63]. Processing statistics are presented in *Supporting information* **Table S1**. The structure of Fab 4G2—PUUV Gc was solved by molecular replacement in Phenix-MR [64] using PUUV Gc (PDB 5J81) and a homology model of the Fab 4G2, generated using the SWISS-MODEL server [65] with PDB 6BPB as the template structure, as search models. Structure refinement was performed by iterative refinement using REFMAC [66] and Phenix [64]. Coot [67] was used for manual rebuilding and MolProbity [68] was used to validate the model. Molecular graphics images were generated using PyMOL (The PyMOL Molecular Graphics System, Version 1.7.0.3, Schrödinger, LLC) and UCSF Chimera [69].

### Preparation of PUUV virus-like particles

PUUV VLPs were produced by transient expression of the complete PUUV M-segment (GenBank CCH22848.1) cloned into the pHLsec vector [55] in HEK293T cells. Six five-layer 875-cm^2^ flasks (Falcon) were used to produce 750 mL of VLP-containing media that was clarified at 3000×g for 20 min to remove cell debris and filtered through a 0.45-μm filter. The virus containing medium was concentrated down to approximately 30 mL using a pump-powered filter (100-kDa cut-off; Vivaflow, Sartorius) and then dialysed into an excess of buffer (10 mM Tris pH 8.0-150 mM NaCl) through a 1-MDa cut-off dialysis membrane (Biotech CE Tubing, Spectrum Chemical) for a few days. The media was further concentrated to ~3 ml with a 100 kDa cut-off centrifugal concentrator filter (Amicon Ultra, Merck Millipore) and layered onto a 20-60% w/v sucrose density gradient in PBS buffer. The gradient was prepared using a Gradient Master (BioComp Instruments, Canada) in a SW32 Beckman tube, and the VLP banded by ultracentrifugation at 4°C for 4 h at 25,000 rpm. The diffuse band (volume of ~3-4 ml) was collected manually, diluted to 20 mL with PBS and pelleted through a cushion of 10% sucrose in PBS to further clean and concentrate the sample (SW32 Beckman centrifuge tube at 25,000 rpm at 4°C for 2 h). Finally, the pellet was resuspended in 60 μl of PBS and stored at 4°C.

### Cryo-EM grid preparation, data acquisition and data processing

A 3-μl aliquot of VLP sample supplemented with 3 μl of 6-nm gold fiducial markers (Aurion) was applied on a holey carbon grid (2-μm hole diameter, C-flat, Protochips) that had been glow discharged in a plasma cleaner (Harrick) for 15 s. The grids were blotted for 3-4 s and plunged into an ethane/propane mixture using a vitrification apparatus (Vitrobot, Thermo Fisher Scientific). For Fab 4G2-treated VLP EM sample preparation, a suspension of purified VLPs was incubated with 2.71 μM Fab 4G2 for 1 h at room temperature prior to grid preparation.

Data were collected using a Titan Krios transmission electron microscope (Thermo Fisher Scientific) operated at 300 kV and at liquid nitrogen temperature. Tomo4 software was used to acquire tomographic tilt data on a direct electron detector (K2 Summit; Gatan) mounted behind an energy filter (0-20 eV; QIF Quantum LS, Gatan). Tilt series were collected from −30 to 60 degrees in 3-degree increments with a dose of 4.5 e^−^ /Å^2^ per tilt and six frames in each movie (*Supporting information,* **Table S2**).

Movie frames were aligned and averaged using Motioncor2 to correct for beam induced motion [70]. Contrast transfer function (CTF) parameters were estimated using CTFFIND4 [71] and a dose-weighting filter was applied to the images according to their accumulated electron dose as described previously [72]. These pre-processing steps were carried out using a custom script named *tomo_preprocess* (available upon request). Tilt images were then aligned using the fiducial markers, corrected for the effects of CTF by phase flipping and used to reconstruct 3D tomograms in IMOD [73]. The amplitudes of the subvolumes cropped from the tomograms were weighted to correct for the low-pass filtering function resulting from dose-weighting of the original images using a custom script (available on request).

Subvolume averaging was performed in Dynamo [74] following previously established procedures [16, 75]. Briefly, initial particle locations (‘seeds’) were created by modelling the VLP membrane surface using the Dynamo’s *tomoview* program. Subvolumes extracted from these locations were iteratively refined against their rotational average, calculated around the membrane model surface normal at each position, to accurately align their locations onto the membrane. Subsequent refinements were carried out using a map of the TULV GP spike (EMDB-4867) as an initial template. The template was low-pass filtered to 50 Å frequency to avoid model bias. Overlapping particles were removed based on a distance filter (106 Å) and cross-correlation threshold after each iteration.

A custom script (PatchFinder; available upon request) was used to divide the spikes to those that were part of a lattice and to those that were not. The script locates the eight closest spikes for each particle, and these are classified as interacting neighbors only if their lattice constraints are within set tolerances (maximum 49-Å and 25-degree deviation from ideal values). Spikes were defined as being part of a lattice if they had at least three interacting neighbors. This classification approach allowed reconstructing the structure of the lattice-bound spike for both VLP datasets (with and without Fab 4G2) in addition to a lattice-free, 4G2-bound spike in the case of the Fab 4G2 sample (*Supporting information*, **Table S3**). In the latter case, the refinement was repeated using 4G2-bound spike as a starting model and by excluding weakly correlating particles (below 0.15). This led to a reconstruction with a lattice-free spike fully decorated with Fab 4G2 around its perimeter (*Supporting information*, **Table S3**). The final maps were filtered to the resolution determined by FSC (0.143 threshold) and rendered as isosurfaces. The surface threshold value was determined according to the molecular weight of the viral ectodomain proteins (and additional Fab) assuming an average protein density of 0.81 Da/Å^3^.

### Quantification of the frequency of lattice formation on VLP surface in the presence and absence of Fab 4G2

The positions of each spike and the number of interacting neighbors were analyzed in PUUV VLPs in the presence and absence of Fab 4G2 from eight VLPs from each cryo-ET dataset. To facilitate the analysis, only VLPs displaying good glycoprotein coverage and roughly spherical morphology were included. Both datasets were processed using the workflow described above, and the PatchFinder script was applied to locate those spikes that were part of a lattice. The particles on the top and bottom of the virus were then removed from further analysis by restricting the second Euler angle (tilt) to 45-135 degrees. This was done in order to exclude those spikes that are facing the air-water interface, which can in itself result in a loss of visible lattice due to its denaturing effect. The number of neighbours for each spike was quantified and the 3D locations of the spikes were plotted in 2D using Mercator projections (*Supporting information*, **Fig S7**).

### Fitting of the crystal structure into the cryo-EM map

The crystal structure of Fab 4G2-PUUV Gc was docked into the reconstruction the Fab-decorated PUUV Gn-Gc spike using the fitmap function in Chimera [69]. A map was simulated for the crystal structure to the resolution of 15 Å, to match the resolution of the spike reconstruction, and fitting was performed using a global search with 500 initial model placements. The top solution (presented in **Fig 5**) yielded a map-to-map correlation score of 0.90 (*Supporting information*, **Table S3**). Gc placement from the top solution was then similarly fitted to into the two reconstructions of PUUV VLP lattice: (i) PUUV VLP treated with 4G2 (reconstruction of latticed areas) and (ii) PUUV VLP (no Fab treatment). Local fitting using the fitmap function in Chimera yielded correlation scores of 0.81 (PUUV VLP treated with Fab 4G2, latticed areas) and 0.82 (PUUV VLP without Fab).

### Accession Codes

Atomic coordinates and structure factors of the PUUV Gc-Fab 4G2 complex, and EM structures of the PUUV VLP surface alone and in complex with Fab 4G2, will be deposited in the Protein Data Bank and the EM Data Bank, respectively, upon final acceptance of the manuscript.

## Acknowledgements

We thank Diamond Light Source for beamtime (proposal MX19946) and the staff of beamline I24 at Diamond Light Source for assistance with data collection. We thank the MRC (MR/L009528/1 and MR/S007555/1 to T.A.B.; MR/N002091/1 to K.J.D. and T.A.B.; MR/K024426/1 to K.J.D.), Academy of Finland (309605 to I.R.) and European Research Council under the European Union’s Horizon 2020 research and innovation programme (649053 to J.T.H.). The Wellcome Centre for Human Genetics is supported by Wellcome Centre grant 203141/Z/16Z. We thank Dimple Karia, Bilal Qureshi, and Adam Costin for electron microscopy support. The Oxford Particle Imaging Centre was founded by a Wellcome Trust JIF award (060208/Z/00/Z) and is supported by equipment grants from WT (093305/Z/10/Z) and MRC. We acknowledge the use of Central Oxford Structural Molecular Imaging Centre (COSMIC) facilities and CSC – IT Center for Science, Finland, for computational resources. Computation also used the Oxford Biomedical Research Computing (BMRC) facility, a joint development between the Wellcome Centre for Human Genetics and the Big Data Institute supported by Health Data Research UK and the NIHR Oxford Biomedical Research Centre. The views expressed are those of the author(s) and not necessarily those of the NHS, the NIHR or the Department of Health.

