## Supporting Information for "Molecular rationale for hantavirus neutralization by a reservoir host-derived monoclonal antibody"

**Items:**

**Figure S1**. Sequence alignment of bank vole (*Myodes glareolus*) Fab 4G2 variable regions with mouse (*Mus musculus*) homologues.

**Figure S2**. Electron density at the Fab 4G2–PUUV Gc interface.

**Figure S3**. Site-directed mutagenesis of Arg100, a centrally located paratope residue, reduces the neutralizing potency of 4G2.

**Figure S4**. Overlay analysis of viral class II fusion proteins with PUUV $Gc_{4G2}$ indicates that PUUV Gc has crystallized in the pre-fusion conformation.

**Figure S5.** Fab 4G2 binding is incompatible with the crystallographically-observed post-fusion conformation of PUUV Gc.

**Figure S6**. The surface of PUUV VLP displays ordered regions of glycoprotein lattice, congruent with previously published reconstructions.

**Figure S7**. Treatment with Fab 4G2 alters the presentation of the hantaviral glycoprotein lattice at the surface of PUUV VLPs.

**Table S1.** Crystallographic data collection and refinement statistics for Fab 4G2–PUUV Gc.

**Table S2.** Cryo-EM tomography data collection statistics.

**Table S3.** Cryo-EM sub-tomogram reconstruction and fitting statistics.

Supplementary References

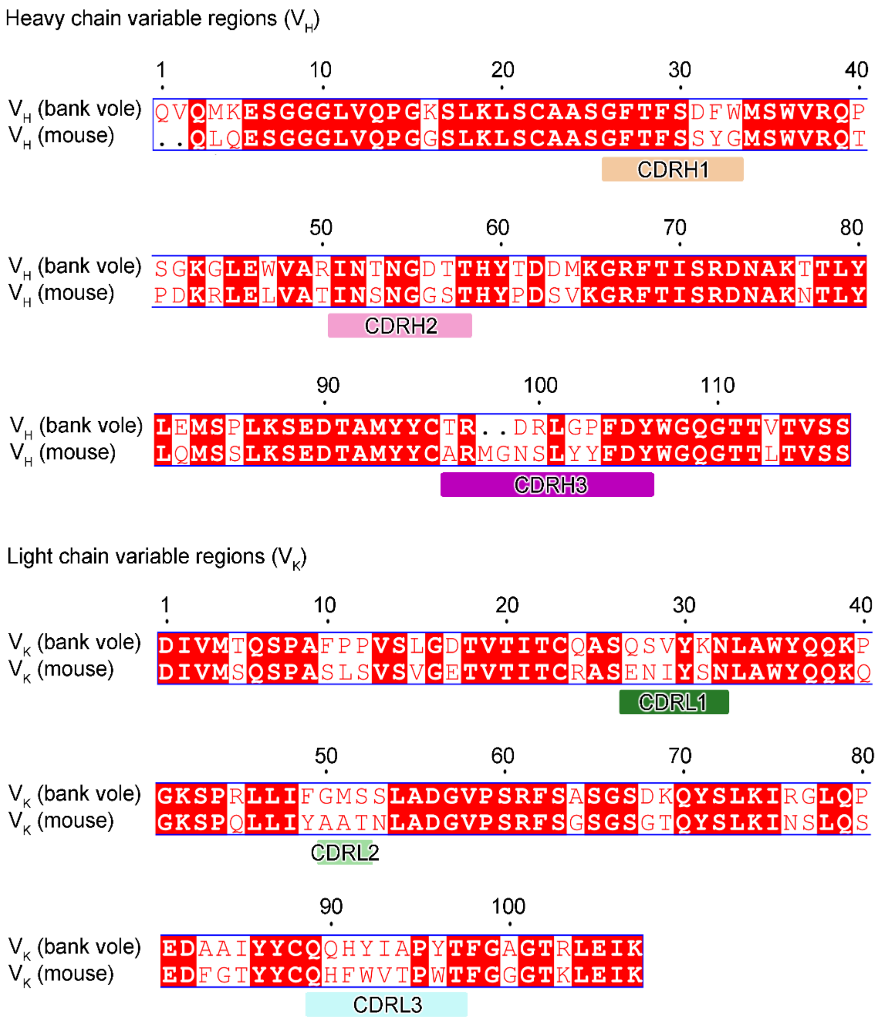


**Fig S1.** Sequence alignment of bank vole (*Myodes glareolus*) 4G2 variable regions with variable regions from representative mouse (*Mus musculus*) antibodies. Mouse homologues used in the alignment were identified by NCBI BLAST [1, 2]. The 4G2 heavy chain variable region (VH) was aligned with the VH sequence from mouse IgG (GenBank: BAN14007.1). The 4G2 light chain variable region (VK) was aligned with the VK sequence from mouse IgG (GenBank: ATI98427.1). Complementarity-determining regions (CDRs) of 4G2 are annotated under the alignment.

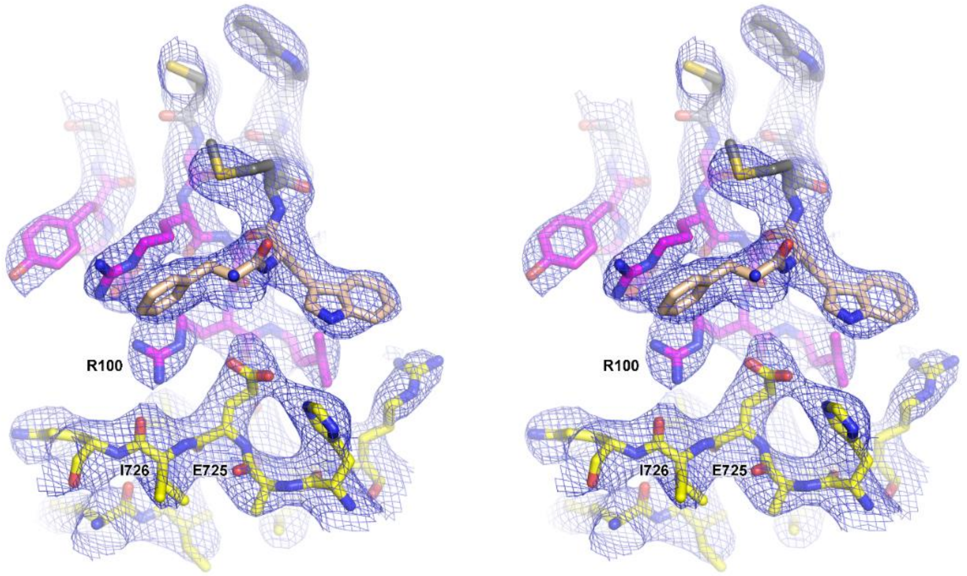


**Fig S2.** Electron density at the Fab 4G2–PUUV Gc interface. Stereo-view of the interaction between PUUV Gc and CDR-H3 of Fab 4G2, with 2*Fo-Fc* electron density map shown contoured at 1σ. The structure is shown in stick representation, with nitrogen atoms colored blue, oxygen atoms red, and carbon atoms yellow for Gc, salmon for Fab 4G2 CDRH2, and magenta for Fab 4G2 CDRH3, respectively. Arginine 100 of CDRH3 and residues Ile726 and Glu725 of Gc, that are predicted by PDBePISA server [3] to form hydrogen bonds and a salt bridge with Arg100, are labelled.

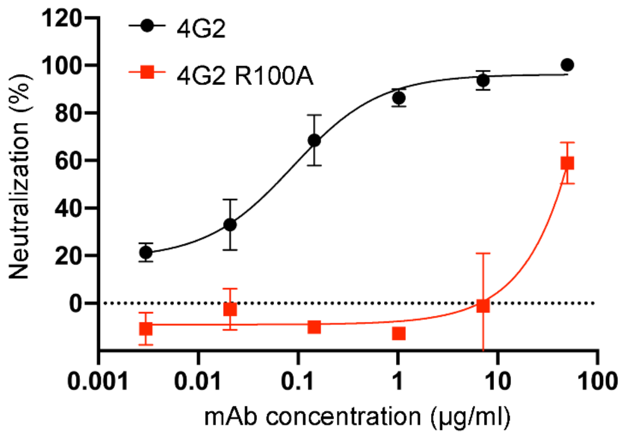


**Fig S3.** Site-directed mutagenesis of Arg100, a centrally located paratope residue in the heavy chain, to Ala reduces the neutralizing potency of mAb 4G2. We compared the neutralization potency of mAb 4G2 R100A mutant (red trace) to that of the wild-type mAb 4G2 (black trace) against PUUV pseudovirus in a hantavirus-pseudotyped VSV-ΔG RFP neutralization assay. Each neutralization assay was carried out three times in duplicate. A representative experiment is shown. Error bars represent the range of the value for the experiment performed in duplicate. A considerable reduction in the neutralization potency of 4G2 R100A was observed, supporting the key role of Arg100 in antigen recognition.

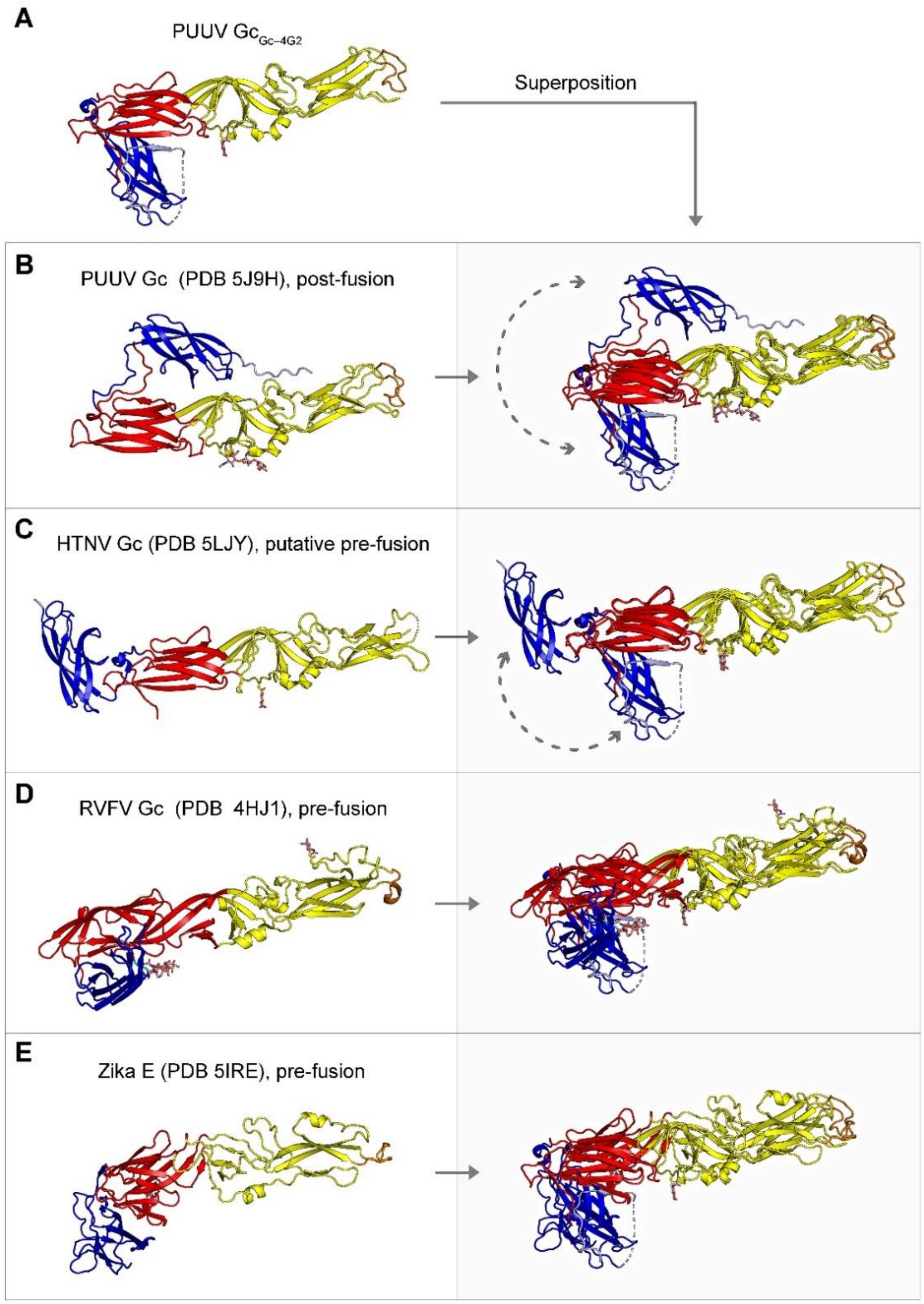


**Fig S4.** Overlay analysis of viral class II fusion proteins with PUUV Gc$_{Gc-4G2}$ indicates that PUUV Gc has crystallized in a conformation similar to the pre-fusion conformation. (**A**) The conformation of PUUV Gc$_{Gc-4G2}$ is distinct from (**B**) the previously reported PUUV Gc post-fusion structure (PDB 5J9H) [4], and aligns more closely with (**C**) the putative pre-fusion configuration of HTNV Gc (PDB 5LJY, assigned as pre-fusion based on dissimilarity from the post-fusion configuration) [5], and (**D**) experimentally confirmed pre-fusion conformations of RVFV Gc (PDB 4HJ1) [6] and (**E**) Zika E (PDB 5IRE) [7]. Cartoon representations of the structure are colored according to domain boundaries (red, yellow and blue for domains I, II and III, respectively). Dotted arrows highlight conformational differences between overlaid structures.

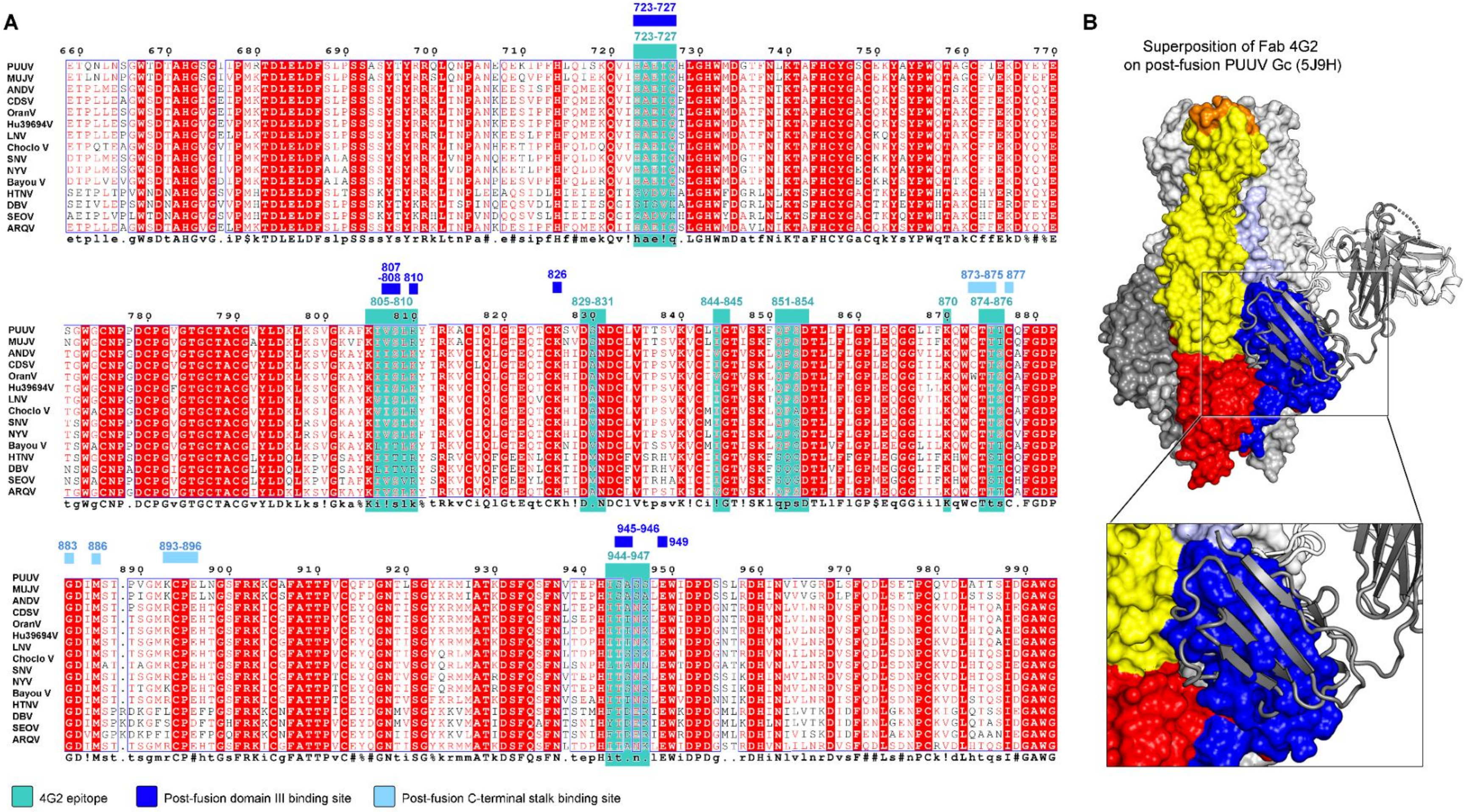


**Fig S5.** Fab 4G2 binding is incompatible with the crystallographically-observed post-fusion conformation of PUUV Gc. (**A**) Sequence alignment of Gc glycoproteins from pathogenic hantaviral species [8, 9], numbered according to PUUV Gc. Residues that comprise the epitope of 4G2 are highlighted in turquoise, and residues that accommodate fusogenic conformational changes are highlighted in shades of blue (dark blue for the binding site of domain III and light blue for the binding site of the C-terminal stalk). In the alignment, fully conserved residues are shown white on red background, partially conserved residues red on white background, and variable residues are rendered black and blue. Hantaviral species included in the alignment comprise Puumala virus (CAB43026.1), Muju virus (MUJV; AGE45110.1), Andes virus (ANDV; AAO86638.1), Sin Nombre virus (SNV; AIA08876.1), New York virus (NYV; AAC54561.1), Laguna Negra virus (LNV, YP_009506658.1), Bayou virus (YP_009505597.1), Choclo virus Gc (APD78411.1), Castelo dos Sonhos virus (CDSV; AAG24914.1), Oran virus (AAB87910.1), Hu39694 virus (AAB87909.1), Araraquara virus (ARQV; AAX78202.1), Hantaan virus (HNTV; AIL25319.1), Seoul virus (SEOV; AXK59723.1) and Dobrava-Belgrade virus (DOBV; AFI98397.1). (**B**) Fab 4G2 binding is incompatible with the post-fusion conformation of Gc. A superposition of Fab 4G2 from the Fab 4G2–PUUV Gc crystal structure onto the crystallographically observed post-fusion conformation of PUUV Gc (PDB 5J9H) [4] reveals that observed epitope is occluded by domain III.

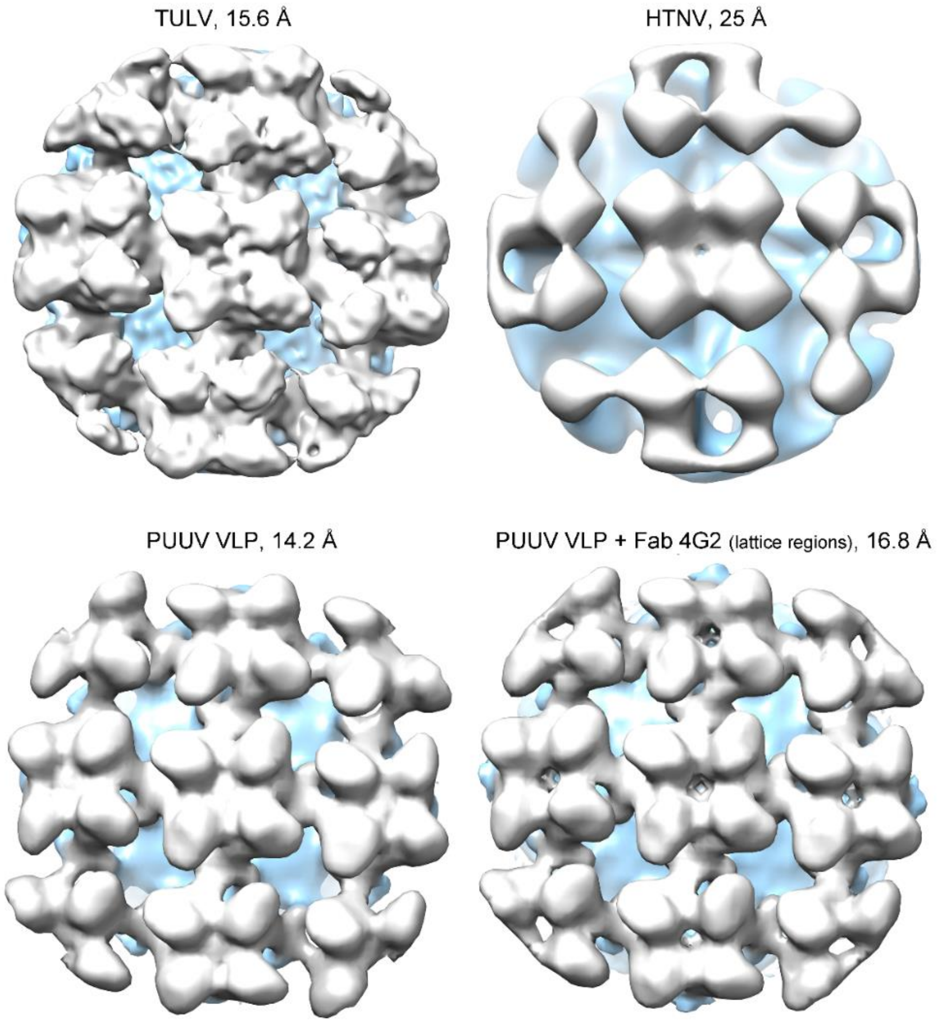


**Fig S6.** The surface of PUUV VLP displays ordered regions of glycoprotein lattice, congruent with previously published reconstructions. Despite the variable resolutions of the present reconstructions of hantaviral surfaces, the tetragonal lattice organization observed in PUUV VLP (14.2 Å) and in PUUV VLP treated with 4G2 (16.8 Å), is similar to that of Tula virus (TULV, 15.6 Å, EMDB-4867) [10] and Hantaan virus (HTNV, 25 Å, EMDB-2056) [11]. Top view of each viral surface reconstruction is shown, and densities corresponding to viral membrane and envelope glycoproteins are rendered in light blue and shades of grey and white, respectively.

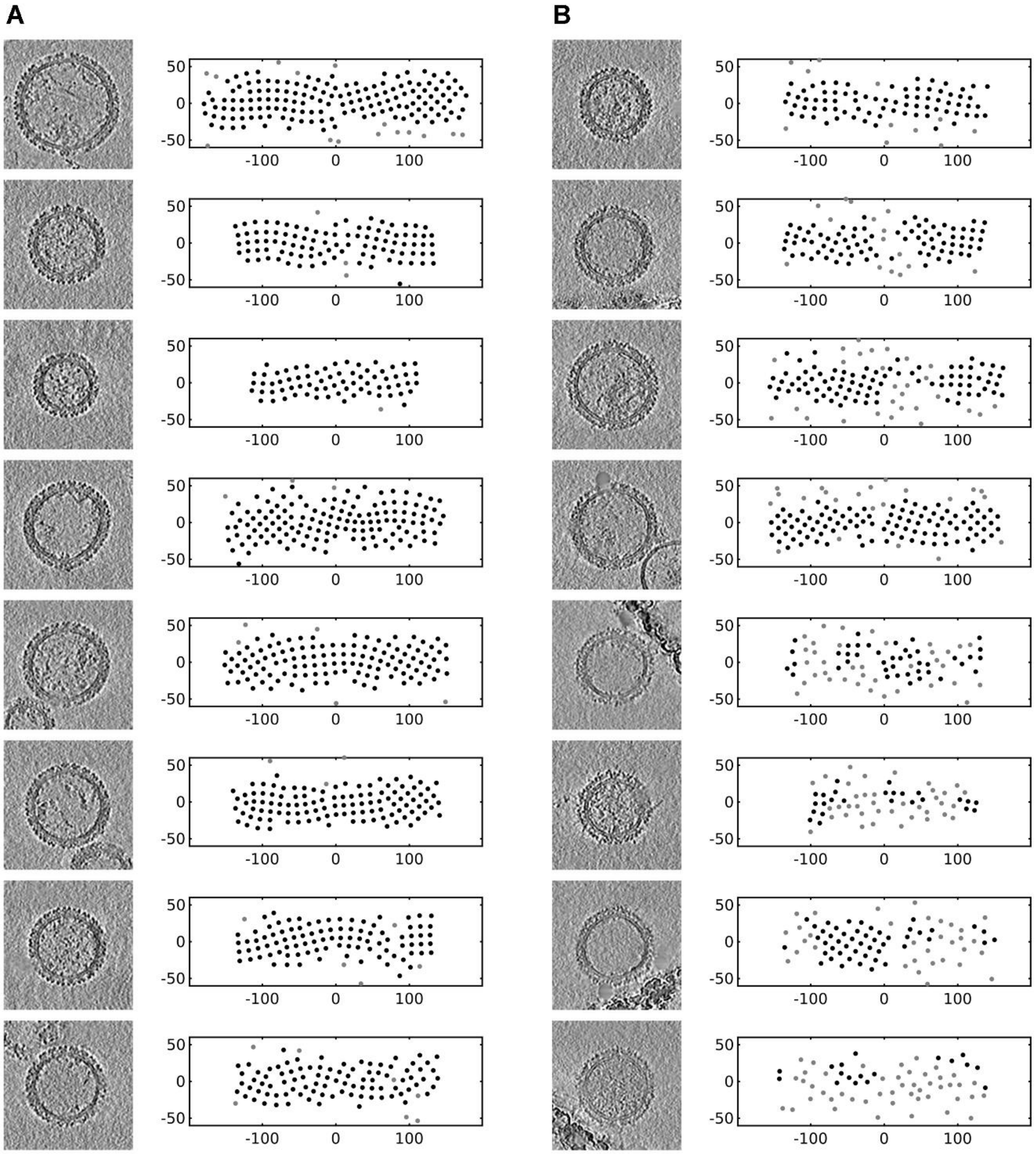


**Fig S7.** Treatment with Fab 4G2 alters the presentation of the hantaviral glycoprotein lattice at the surface of PUUV VLPs. Tomographic slices of the selected PUUV VLPs (left) and corresponding Mercator projections [12] (right) describing the positions of $(Gn–Gc)_4$ spikes on VLP surfaces are shown (**A**) in the absence and (**B**) presence of Fab 4G2. The subset of spike positions that belong to regular patches (determined using the PatchFinder script; please see main Methods) are displayed in black. Units shown are in nanometers.

**Table S1.** Crystallographic data collection and refinement statistics for Fab 4G2–PUUV Gc.

| | Fab 4G2–PUUV Gc |
|---|---|
| **Data collection** | |
| Space group | *P* 1 21 1 |
| Cell dimensions | |
| *a*, *b*, *c* (Å) | 97.2, 50.5, 124.5 |
| *a*, *b*, *g* (°) | 90.0, 107.7, 90.0 |
| Resolution (Å) | 92.6-3.50 (3.63-3.50)* |
| $R_{merge}$ | 0.413 (-) |
| $R_{pim}$ | 0.140 (0.426) |
| I/σ I | 4.6 (1.9) |
| $CC_{1/2}$ | 0.96 (0.65) |
| Completeness (%) | 99.7 (97.5) |
| Multiplicity | 9.6 (9.0) |
| **Refinement** | |
| Resolution (Å) | 92.63-3.50 (3.77-3.50) |
| No. reflections | 14,903 |
| $R_{work}$ / $R_{free}$ | 0.214 / 0.263 |
| No. atoms | |
| Protein | 6,467 |
| Ligand/ion | 32 |
| Water | 0 |
| *B*-factors | |
| Protein | 69.8 |
| Ligand/ion | 58.3 |
| Ramachandran plot (%) | |
| Favored region | 91.16 |
| Allowed region | 8.84 |
| Outliers | 0 |
| R.m.s deviations | |
| Bond lengths (Å) | 0.001 |
| Bond angles (°) | 0.412 |

* Highest resolution shell is shown in parentheses

**Table S2.** Cryo-EM tomography data collection statistics.

| | **PUUV VLP** | **PUUV VLP + 4G2 fab** |
|---|---|---|
| Tilt range (degrees) | –30–60 | –30–60 |
| Tilt increment (degrees) | 3 | 3 |
| Frames per tilt | 6 | 6 |
| Electron exposure ($e^-$/ Å$^2$/tilt) | 4.5 | 4.5 |
| Total electron exposure ($e^-$/ Å$^2$) | 140 | 140 |
| Pixel size (Å) | 1.77 | 1.76 |
| Defocus range (μm) | 2.8–4.0 | 2.8–4.0 |
| Tilt series | 33 | 59 |
| Tomograms | 33 | 59 |
| VLP sub-volumes | 42 | 79 |

**Table S3.** Cryo-EM sub-tomogram reconstruction and fitting statistics.

| | PUUV VLP | PUUV VLP + 4G2 fab | |
|---|---|---|---|
| Reconstruction | GP spike | GP spike | GP spike+4G2 |
| EMDB ID | EMD- xxxx | EMD- xxxx | EMD- xxxx |
| Spike sub-volumes | 4,333 | 1,998 | 1,721 |
| Symmetry | C4 | C4 | C4 |
| Pixel size (Å) | 7.08 | 7.04 | 7.04 |
| Resolution (Å) | 14.2 | 16.8 | 15.0 |
| Model-to-Map CC | 0.82^ | 0.81^ | 0.90* |

CC: cross-correlation from Chimera fitmap
* Fitted structure: Fab 4G2–PUUV Gc complex crystal structure
^ Fitted structure: PUUV $Gc_{Gc–4G}$
